# Collagen-Based Gut-on-Chip for *in vitro* modeling of intestinal barrier function and host-pathogen interactions

**DOI:** 10.1101/2025.05.24.655930

**Authors:** Fernanda López García, Jacques Leng, Hugo de Oliveira, Pierre Nassoy, Karine Dementhon

## Abstract

The human intestine, characterized by its villi and crypt structures, plays a critical role in nutrient absorption, barrier function, and host defense. However, traditional *in-vitro* models employing synthetic membranes like polydimethylsiloxane (PDMS) and polycarbonate (PC) often fail to accurately replicate the complex physiological environment of the intestine. To address this limitation, we developed a gut-on-chip, incorporating a porous collagen type I membrane, to better mimic the natural extracellular matrix (ECM) and create a more physiologically relevant *in vitro* system. Thin, porous collagen type I membranes were fabricated and characterized by linear close contact profilometry to determine their thickness, which closely approximated the *in vivo* intestinal basement membrane. Caco-2 cells cultured within the device exhibited the formation of villi-like structures, tight junction formation, and mucin production, demonstrating successful differentiation and functional barrier formation on the collagen membrane. We investigated the device’s capacity to model host-pathogen interactions by infecting the cell layer with *Candida albicans*. Confocal microscopy revealed hyphal invasion of the epithelial cells, and permeability assays demonstrated increased layer permeability following infection, highlighting the device’s ability to replicate infection processes and their impact on barrier integrity. This gut-on-chip, by integrating physiological membrane and replicating key structural and functional aspects of the intestine, offers a promising platform for studying intestinal physiology and host-pathogen dynamics.

## Introduction

The human intestine plays an important role in digestion, absorption of nutrients, and protection of the human body (1). All of these functions are largely mediated by the inner surface layer of epithelial cells in the intestine with its distinctive finger-like protrusions known as villi and deep cavities termed crypts (2). Villi, formed by circular folds of mucosa and submucosa, serve to increase the absorptive surface area of the tissue, while crypts house proliferative stem cells that differentiate and migrate upwards toward the villus tips (3,4). Another major feature of the intestine environment is the presence of mechanical forces essential for luminal content transport, including peristaltic movements driven by smooth muscle contractions and shear stress induced by luminal fluid flow (5). These mechanical cues stimuli play critical roles in regulating epithelial cell differentiation, proliferation, and barrier function (6,7).

Organ-on-chip (OOC) microfluidic technology offers a promising approach to recapitulate the architectural and functional features of the intestine and to assess host-pathogen interactions. In recent years, microfluidic systems have experienced significant advancements in biomedical and biological research, driven by their rapid and straightforward fabrication, customizable designs, and minimal requirements for reagents and biological samples (8). Two principal strategies have been established for the development of gut-on-a-chip platforms (9,10). The first one consists in the fabrication of molds or scaffolds that recapitulate crypts and villi and, which are subsequently coated with intestinal cells (11-13). Engineering these architectural cues not only mimics intestinal structure but also proved successfully to capture key functional aspects of intestinal physiology (14). The second one starts with a flat surface, i.e. a porous membrane that separates two channels within the microfluidic device. This design emerges from the initial developments pioneered by Ingber and colleagues (8,15,16). In each compartment, two or more different cell lines can coexist, allowing communication and exchange of cues and nutrients (17). Physiomimetism relies on cell self-organization instead of scaffolding. 3D micro-architecture in crypt-and villi-like structures has been shown to emerge spontaneously if shear is applied (18). However, the nature of the material used to fabricate the porous membrane seems to be crucial. Perforated polydimethylsiloxane (PDMS) sheets are commonly used. Their advantages are transparency, flexibility, biocompatibility and capacity to replicate intricate designs. Polycarbonate (PC) is another popular material used due to its commercial availability and low prices (19). Despite their biocompatibility, these materials have two main drawbacks. The stiffness of the membranes has been sometimes reported to be sub-optimal for spontaneous morphogenesis (20). Moreover, a surface treatment is necessary to control the hydrophilicity and functionalize the surface with natural polymers such as ECM proteins (collagen, fibronectin, etc.) to ensure cell adhesion (21).

Given the limitations of synthetic materials such as PDMS and PC, an alternative approach is to consider the extracellular matrix (ECM) as a potential membrane material. In the intestine, ECM is formed by a network of fibrous structural proteins that act as a scaffold to hold together the representative intestinal architecture, and proteoglycans that fill the space between fibers in the form of a gel. The intestinal ECM is divided into two layers: the basement membrane (BM) and the interstitial membrane (IM). The BM is produced by both the epithelial and mesenchymal cells, and its main function is to help regulate epithelial cells homeostasis and provide structural support for the epithelium (22). The major structural component of the BM is collagen type IV that forms open non-fibrillar networks (23). Beneath the BM lies the IM, which is mainly composed of fibrillar collagen, including types I and III (24). Collagen type I is the most abundant in the intestinal interstitial matrix, providing tensile strength and structural support through dense, organized fibers, while collagen type III contributes to elasticity and flexibility, helping maintain tissue homeostasis with its more loosely associated and pliable structure (24). Various groups have actively investigated the potential use of ECM-derived membranes as inserts for cell culture or OOC. These include vitrified membranes made from purified components such as collagen (21, 25, 26), fibronectin, or laminin (27), as well as complex mixtures such as Matrigel and decellularized matrix (28). However, their difficulty to handle and inherent lack of mechanical stability was reported to be a limitation for their use in long-term and dynamic cell or tissue culture applications. Consequently, these materials were reinforced by combining them with nanofibers such as PCL, silk fibroin (19, 29, 30) or made stretchable by adding elastin (31).

In this work, we present the development of a gut-on-chip platform by employing a native porous collagen type I membrane, strategically chosen for its fibrillar structure. This approach seeks to transcend the limitations of synthetic membranes, and thereby more accurately replicate the *in vivo* environment of the human intestine. The morphological and mechanical properties of the fabricated collagen I membranes were characterized in depth. Following the assembly of the 2-chamber microfluidic device separated by the collagen membrane, Caco-2 cells were seeded within the upper channel. We showed that continuous fluid flow, without peristaltism induced by cyclic deformation, as suggested in the pioneering works that used a synthetic membrane, promotes rapid cellular differentiation of the cell monolayer. This differentiation was confirmed by key markers of tight junction formation, mucus production, by the recapitulation of the three-dimensional crypt-villi structure using scanning electron microscopy and confocal microscopy, and permeability measurements demonstrating the barrier function of the epithelium. Finally, the differentiated epithelial layer was challenged with infection by *Candida albicans* yeast which was shown to disrupt the barrier function through hyphae penetration into the intestinal epithelium, and even damage to the collagen membrane.

## Materials and methods

### Collagen type I membranes fabrication

Collagen type I membranes were fabricated by adapting a previously described protocol (22).The protocol is sketched in Fig. 1A. Briefly, a volume of 800 µl (for each membrane) of a solution of bovine collagen type I (TeloCol ^®^10, Cellsystems, Germany) at a 2 mg/ml concentration in Dulbecco’s Modified Eagle Medium Glutamax (DMEM; Gibco, France) supplemented with 10% decomplemented fetal bovine serum (FBS; Gibco) with antibiotics (50 µg/ml of penicillin and streptomycin) (further referred to as cDMEM) was spread on a polydimethy-siloxane (PDMS; Sylgard 184, Dow Corning) surface. After gelation by incubation at 37°C for 1 hour, the membranes were left to dry overnight in sterile conditions in a laminar flow hood. Then, the membranes were washed with sterile deionized water and rehydrated for 4 hours. The excess of water was aspirated, and the membranes were left to dry overnight. The membranes could then be peeled off easily from the PDMS surface and manipulated using tweezers without any risk of damage. Finally, they were kept in a dry petri dish until use.

**Figure 1:**
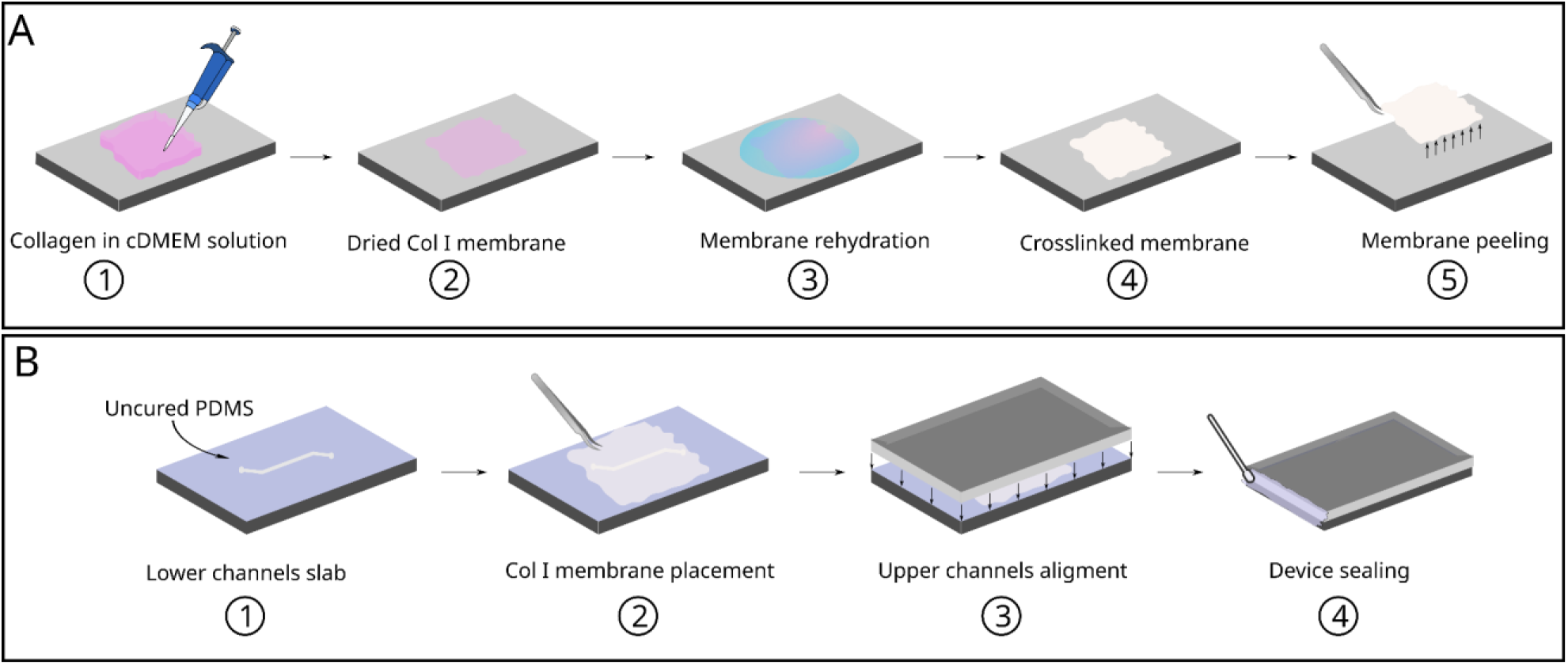
Chip assembly with a collagen membrane. **A. Fabrication of collagen type I membranes.** 1) Collagen I in cDMEM solution is spread onto PDMS surface; 2) Drying by evaporation of the solution at room temperature; 3) Membrane washing with sterile water to get rid of culture media crystals and phenol red; 4) Drying by evaporation of collagen membrane at room temperature; 5) The membrane is peeled off from PDMS surface. **B. Assembly of the microfluidic device**. 1)Lower channel PDMS slab covered with 5:1 PDMS prepolymer; 2) Positioning of the collagen membrane over the lower channels; 3) Alignment and positioning of upper channels slab covered with PDMS prepolymer and 4) Sealing of the device with PDMS prepolymer to avoid leakage.

### SU-8 mold fabrication

Photoresist (SU-8 2050, Microchem, Newton, MA) was spin-coated over a silicon wafer using a 1500 rpm speed. The photoresist layer was heated at 95°C for 5 min and cooled down to room temperature. To reach a 250 µm thickness for the lower channels and 750 µm for the upper channels, the operation was repeated as many times as required (2 times for the lower channels and 4 times for the upper channels). A final soft-baking of 15 min at 95°C was performed to evaporate the remaining solvent. The coated wafer then underwent a discontinuous UV light exposure process (Süss Microtec, MJB4 mask aligner), consisting of 30 seconds of UV light followed by a 3-second pause, repeating this cycle 3 times. A post-exposition baking was performed at 95°C for 1 hour and the wafer was then transferred to a glass beaker with developer solution for 15 min in a shaker to reveal the pattern. The wafer was rinsed with ethanol and dried on a heating plate at 95°C for 10 min.

### Chips fabrication and assembly

The device was fabricated by casting PDMS (Sylgard 184, Dow Corning) with a base: curing agent ratio of 10:1 (w/w) on the molds with the negative designs of the upper and lower channels made on silicon wafers with SU-8 photoresist. The prepolymer was put in the oven at 65°C for 1 hour. After crosslinking, upper and lower channels-containing slabs were cut to the desired dimensions. To assemble both pieces with the collagen membrane sandwiched in between, we followed the same strategy as developed in (21). The protocol is shown in Fig. 1-B. A prepolymer of PDMS 5:1 (w:w) was prepared and spin-coated on a Petri dish. Then, the lower PDMS slab was dipped into the layer of prepolymer making sure the channels were not covered by the prepolymer and clogged. Within 5 min, the collagen membrane was placed on top of the PDMS slab and it was cured overnight at room temperature. Finally, the top PDMS slab was dipped into the 5:1 prepolymer and placed on top of the collagen membrane making sure the upper and lower channels were aligned and taking out the air bubbles that may be trapped between the layers. The assembly was completed by curing overnight at room temperature.

### Scanning electron microscopy (SEM)

For SEM imaging, Collagen type I membranes were fixed using 2.5% glutaraldehyde and 2% paraformaldehyde (PFA) in PBS at room temperature overnight. They were then washed 3 times with PBS and dehydrated in a graded ethanol series (30%, 50%, 70%, 80%, 90%) with a 10 min incubation with each concentration and a final dehydration in two 10-minute incubations with 100% ethanol. After air drying at room temperature overnight, membranes were metalized with a layer of gold. Micrographs were obtained on a tabletop scanning electron microscope (TM4000, Hitachi) and analyzed using ImageJ software to measure the average pore size.

To observe the cells seeded inside the microfluidic by SEM, cells were fixed by running 4% PFA inside upper and lower microchannels for 2 hours. The channels were separated from each other to make the cells more accessible. Cells were washed 3 times with PBS before being dehydrated by successive 15-min incubations in graded 50%,70%,80%, 90% ethanol baths, followed by two 15 min final baths with 100% ethanol. The cells were left to dry overnight at room temperature. As a final dehydration step, the cells were incubated twice for 10 minutes in hexamethyldisilazane (HMDS). Finally, samples were obtained by cutting the chip with a biopsy puncher of 6mm and metalized with a layer of gold prior to observation by SEM (TM4000, Hitachi).

### Profilometry

The average thickness of collagen membranes was obtained using a contact profilometer (Veeco Instruments, Dektak 6M). To assess the reproducibility of our protocol, different collagen I membranes were measured (n=6). The membrane was electrostatically bonded to a microscope glass slide. A stylus with a 12.5 µm radius was used on the close contact setting of the profilometer; the stylus force established for the measurement was 3 mg along 3000 µm of the membrane surface. In order to have a reference height the stylus point was made to trace a line going from the membrane to the glass slide surface (representing the zero).

### Tension test on collagen membranes

For testing the mechanical properties of dry collagen membranes, three membranes were fabricated using the protocol described in the previous section in a dog-bone shape rectangle. A TA-HD-Plus texture analyzer (Stable Micro Systems) with 5 kg load cell was used to perform these tests. The membranes were placed vertically between the two grips of the apparatus and adjusted until tense. Then the load was applied until failure and the data was registered, tests were done on triplicate for each membrane. Each membrane width was measured to derive its transversal area. Finally, the tensile stress (in MPa) is given by: *σ*=F/A, where F is the applied force in Newtons and A is the transversal area of the membranes in mm^2^.

### Cell culture

A Caco-2 cell line (HTB-37), derived from a human colorectal adenocarcinoma (ATCC, Manassas, VA, USA), was cultured and maintained in cDMEM, at 37°C in 5% CO_2_.

### Cell seeding inside microfluidic device

For cell seeding inside the microfluidic device, the upper and lower channels were flushed 2 times with 70% ethanol for sterilization. Then both channels were filled with cDMEM and incubated for 1 hour at 37°C and 5% CO_2_. Caco-2 cells at a 80% confluence were detached from a 75 cm^2^ using 0.25% trypsin EDTA (Gibco, France). Cells were counted, centrifuged and diluted in culture media at a final concentration of 20×10^3^ cells per ml. Cells were then seeded at a density of 2.5×10^6^ cells per cm^2^ cells per cm^2^ in the microfluidic chip. The device was incubated for 4 hours at 37°C and 5% CO_2_ to allow cell adhesion to the membrane. cDMEM was continuously perfused through both channels at constant rates of 8.9 μl/min in the upper channel and 1.3 µl/min in the lower channel, using syringe pumps (Centoni, Nemesys) to achieve a shear stress of 0.02 dynes/cm^2^ on both sides of the membrane.

### Yeast strain and intestinal epithelium infection

*C. albicans* wild-type strain SC5314 was grown in standard YPD (1% yeast extract, 2% bactopeptone and 2% glucose) agar plates at 37°C. Yeast were grown overnight in 5 ml of YPD at 30°C and 200 rpm reaching exponential growth phase. Prior to the infection of the intestinal epithelium yeast cells were counted using a Malassez chamber and were taken to centrifuged at 1000 rpm for 5 min to adjust the concentration at 1.5×10^4^ cells per ml using Roswell Park Memorial Institute 1640 (RPMI; Sigma Aldrich, France) culture media without phenol red and supplemented with 10% decomplemented FBS and antibiotics (50 µg/ml penicillin and streptomycin) (further referred to as cRPMI-PR). On day 7 of Caco-2 cell culture inside the microfluidic device, 50 µl of the yeast suspension was added to the upper channel and incubated for 10 min in static conditions to allow the yeast to adhere to the epithelial layer. Subsequently, the flow was restarted at the same flow conditions as used before. After 6 hours, the infection was stopped by fixation of the cells.

### Immunostaining and fluorescence microscopy

Following the cell culture experiments, the microfluidic devices were disconnected from the pump and the remaining culture media inside the microchannels was removed. Cells on the surface of the collagen membrane were fixed by incubating them with 4% PFA in cytoskeleton stabilization buffer (10 mM 2-Morpholinoethanesulphonic acid (MES), 150 mM NaCl, 5 mM EDTA, 5 mM MgCl_2_, 5 mM glucose, pH 6.1), for 2 hours at room temperature. Then the channels were washed 3 times with PBS to continue with permeabilization and blocking by adding 0.1% Triton-X in PBS for 30 min at room temperature and 10% FBS in PBS for 2 hours at room temperature. Once cells were fixed and blocked, they were incubated with the primary antibodies dissolved in the blocking solution [anti ZO-1 rabbit (1:100), anti MUC-2 mouse (1:50), anti Villin rabbit (1:100)] at 4° C overnight. After three PBS washes, the cells were incubated with the secondary antibodies [anti-rabbit Alexa 488 (1:100), anti-mouse FITC (1:100), anti-rabbit Alexa 594 (1:100)] for 2 hours at room temperature in dark. Finally, the channels were washed 3 times with PBS, and cells were counterstained with DAPI. Yeast cells were stained using Calcofluor White (Sigma-Aldrich, France) at a concentration of 250 ng/µl by incubation for 15 min and washed 3 times with PBS. To obtain fluorescent images we use an inverted confocal microscope (TCS SP5, Leica) with a 25x water lens that allowed us to have a view on the full width of the central channel and to avoid the need to open the chip for cell observation.

### Permeability assays

Permeability assays were performed inside the microfluidic chip in 3 different conditions: naked collagen membrane, differentiated epithelium and finally infected cell layer. To assess the baseline permeability of the collagen membrane, an empty microfluidic device was incubated with cRPMI-PR in both the upper and lower channels at 37° C and 5% CO_2_ for one hour. The upper channel of the microfluidic chip was connected to a syringe prefilled with a 50 µM solution of fluorescein in cRPMI-PR cell culture media and the lower channel was connected to a syringe prefilled with cRPMI-PR, the culture media was perfused in both channels at flow rate of 4 µl/min. At each outlet, an Eppendorf tube was placed as a collection reservoir containing 1080 µL of cRPMI-PR cell culture media to achieve a 10-fold dilution of the fluorescent marker. Samples of 120 µL were collected from each reservoir in triplicate every 30 minutes, to maintain a constant volume, 360 µL of fresh cRPMI-PR medium were added back to the reservoir after each sampling. Samples were deposited in a 96-well plate and fluorescein fluorescence intensity (excitation/emission: 495/520 nm) was measured using a FluoStar Optima fluorometer (BMG Labtech). The results were recorded in arbitrary units of fluorescence (AUF). Corresponding concentration of fluorescein was established using a calibration curve performed using a concentration of fluorescein ranging from 5 µM to 4.8 nM. For the microfluidic devices with healthy differentiated epithelial layer (day 7 of cell culture) and the infected layer (6 hours infection), prior to performing the permeability assay cDMEM culture media was replaced with cRPMI-PR medium, which was perfused during one hour, following the protocol described earlier. The apparent permeability coefficient for each case was calculated according to the provided equation.

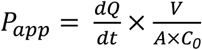

where dQ/dt is the change in concentration the lower channel of the microfluidic device over time, V is the volume of the lower channel (0.0405 cm^3^), A is the surface area of the membrane (0.15 cm^2^) and C_o_ is the initial concentration of fluorescein in the upper channel (50 µM).

## Results and discussion

### Topographical features and mechanical properties of Collagen type I porous membranes

The ECM not only plays a pivotal role in regulating cell morphogenesis and differentiation but also serves as the structural foundation for the intestinal epithelium (33). One of the primary components contributing to this structure and function is collagen that exists in different types (34). To build a stable membrane, we chose Collagen I, the fibrillar structure of which could support cell growth in the chip. We generated thin films from a commercial bovine skin collagen I solution that look like smooth membranes at first sight. However, scanning electron micrographs reveal a more complex surface organization with pores and entangled fibers. After image analysis, collagen fibers are between 500 nm and 1 µm in diameter (figure 2A). Pores observed between the intertwined collagen fibers were derived from image analysis following a standard method described in (35). We found an average diameter of 821 ± 372 nm (Figure 2B). Next, the thickness of the membrane was derived from close contact profilometry experiments. The average thickness of the membrane was found to be 3.75 ± 0.85 µm (figure 2C) Finally, their mechanical properties were assessed under dry conditions at room temperature. Our home-made collagen membrane exhibited an average tensile strength of 83.4 ± 11.9 MPa, indicating a satisfactory reproducibility from the manual manufacture of one membrane to another (figure 2D). By contrast, although the properties of native membranes vary depending on the tissue, they are typically characterized by fibrils of the order of 100 nm in diameter (29), suggesting that collagen fibers bundling events might be enhanced by the drying process of our collagen membrane. Moreover, the thickness and pore size of native membranes were reported to be rather in the 50-100 nm and 10-100 nm respectively (28). The tensile strength of BM is finally in the 1-10 MPa range (28). Relative to the same thickness, the resistance or rupture force of our collagen membrane is therefore equivalent to the one of the BM. Although they are an order of magnitude larger than the native BM pores, the collagen membrane pores should prevent transmigration of most cells. In the same line, the thickness is more than 30 times larger. A lower thickness would be desirable in order to favor the physical contact between cells in both compartments. However, this collagen membrane is already quite an improvement compared to synthetic PDMS membranes, for which the thickness is about 30 µm (36), the pore size 7-10 µm (37) and the tensile strength of the order of 1 MPa.

**Figure 2:**
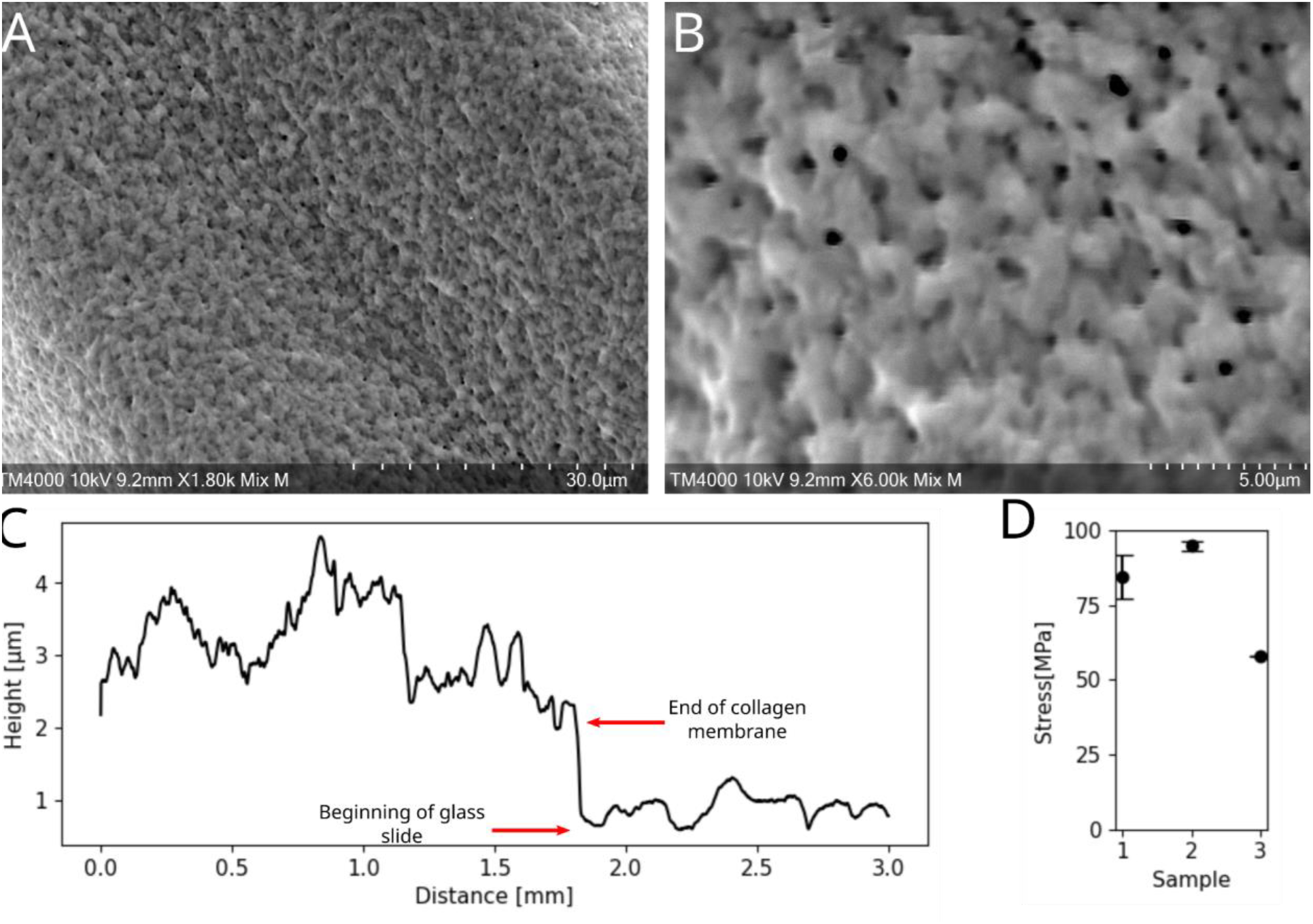
Collagen type I membrane surface and mechanical characterization. A. Low magnification SEM image of collagen type I membrane (scale bar = 30 µm). B. High magnification SEM image of collagen type I membrane (scale bars = 5 µm). C. Representative graph of thickness measurement of 6 collagen membranes by close contact profilometry with an average thickness of 3.7 ± 0.8 µm and D. Tensile test made on dry membranes having an average stress value of 83 ± 12 MPa.

The assembly of the microfluidic device involves several critical steps to ensure proper operation. The first and most crucial step is placing the collagen membrane on the PDMS slab of the lower channel. The membrane was carefully positioned to ensure that the inlet and outlet of the lower channel remain uncovered. Next, the upper channel was aligned with the lower one, ensuring that the both PDMS slabs were in complete contact to prevent any leaks when flow is applied. Figure 3A illustrates the fully assembled microfluidic device, while Figure 3B provides a cross-sectional view of the device. The upper channel was designed to house the intestinal epithelial cells, while the lower channel supplies nutrients to the cells from the basolateral side, as shown in Figure 3C. Figure 3D depicts the entire experimental setup. A computer controls the syringe pumps, which inject the culture medium at a constant flow rate to achieve a shear stress value of 0.02 dyne/cm^2^, mimicking the conditions cells experience in vivo. This shear stress is applied uniformly in both channels. The chip is enclosed in an incubator maintained at 37°C and 5% CO_2_

**Figure 3.**
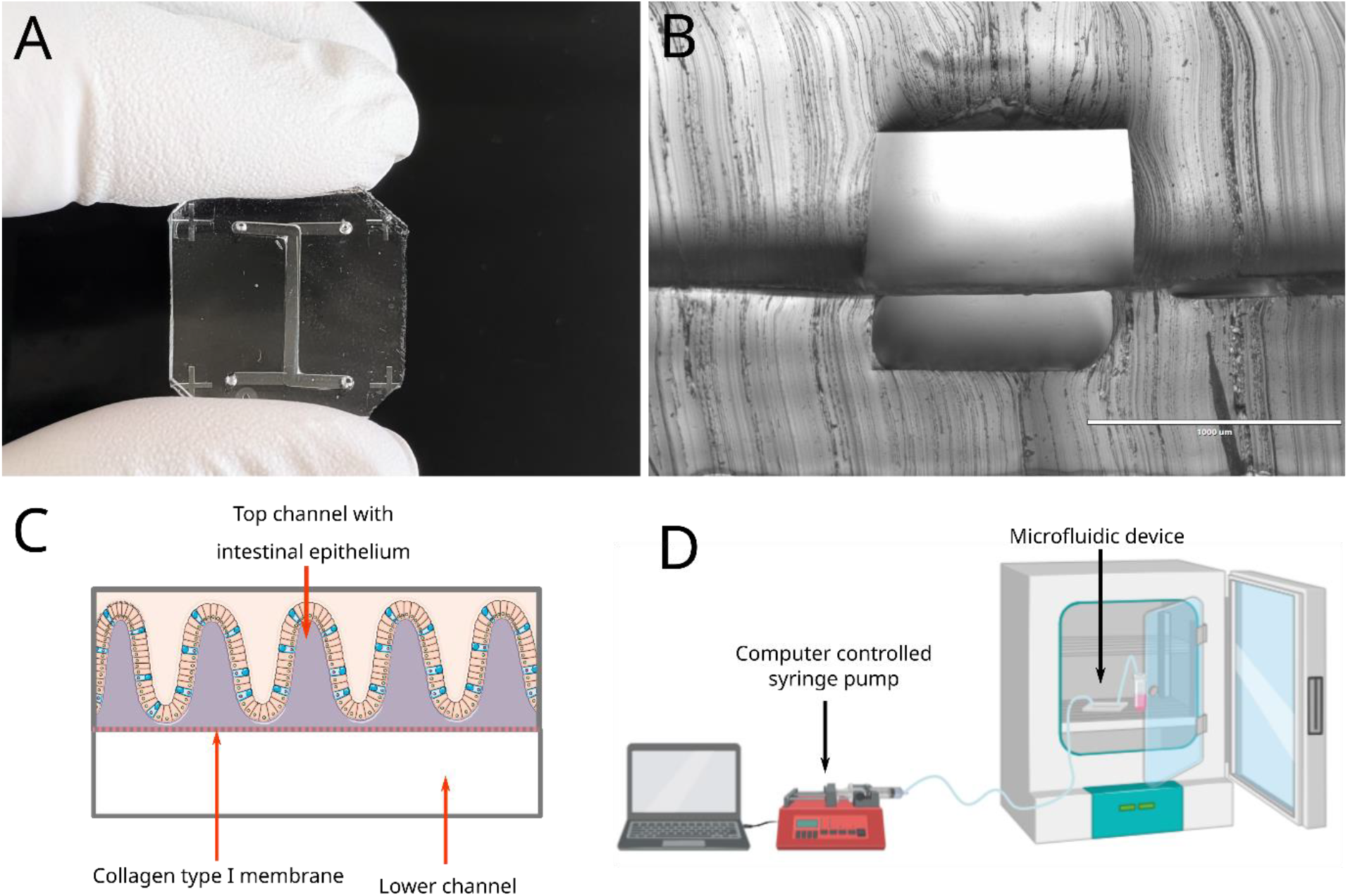
Microfluidic device for intestine on chip model. A. Image of the microfluidic device for cell culture of caco-2 cells. B. Transversal cut of assembled microfluidic device with collagen I membrane. C. Diagram of transversal cut of the microfluidic device with a differentiated epithelial layer. D. Experimental set-up for intestine-on-chip.

### Epithelial cell differentiation and villi-like formation

Caco-2 cells were seeded inside two microfluidic chips to compare the cellular growth and differentiation process during seven days. One chip was maintained under static conditions, while the other was subjected to continuous flow. The influence of constant flow on Caco-2 cell morphogenesis was evident as early as day 2, with the initiation of the cell monolayer showing folds in the device subjected to constant flow. This development occurred prematurely compared to literature reports on similar systems, which typically describe fold formation beginning between days 4 and 5 post-seeding (39). In contrast, cells under static conditions formed a confluent but consistently flat monolayer at days 2 and 4. By day 7, the flow-induced undulations and folds had expanded to cover the entire collagen membrane surface, while the static device only showed the initial signs of these structures. The progression of growth and cell morphogenesis during this time under both conditions is presented in Figure 4.

**Figure 4.**
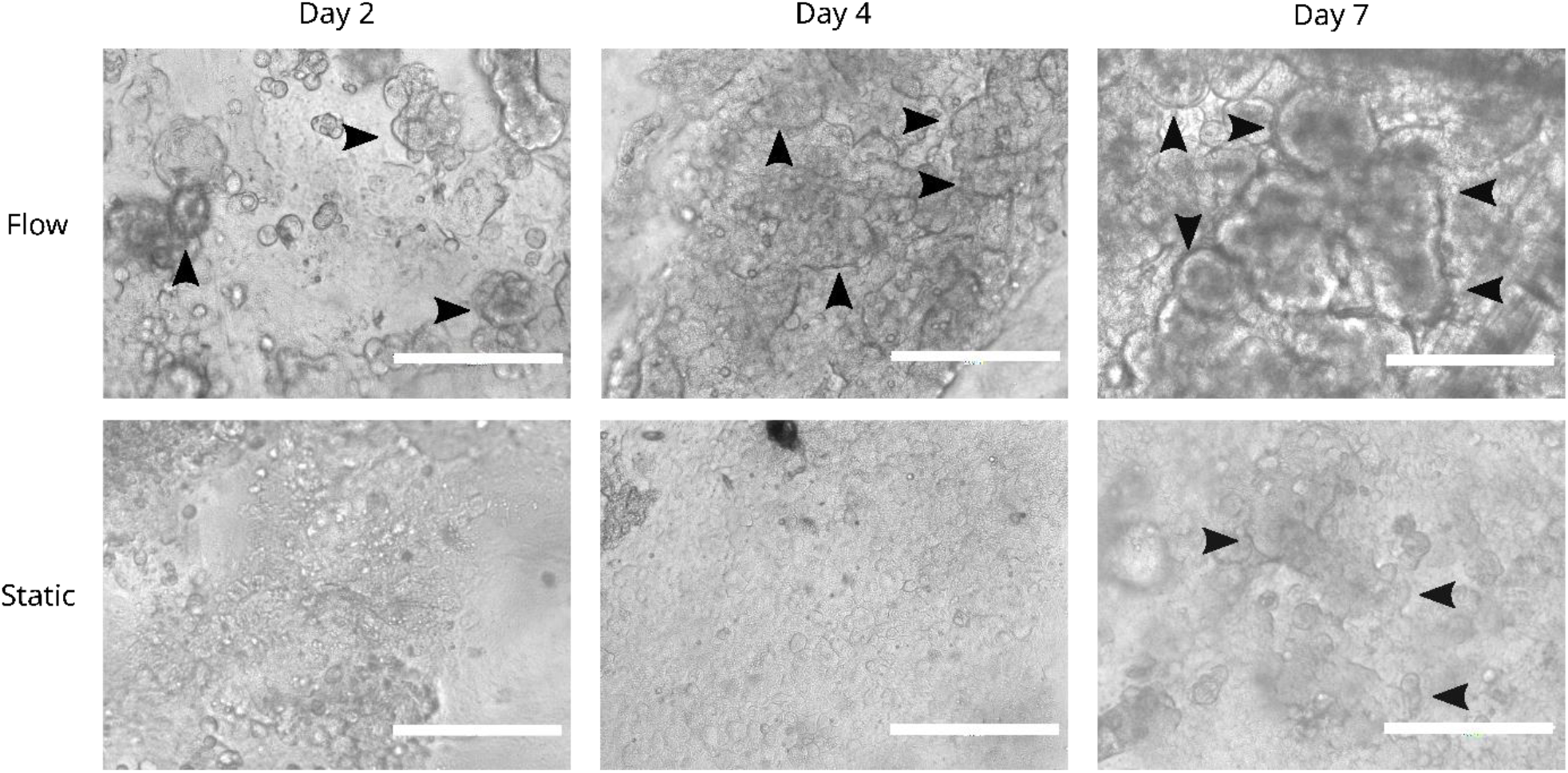
Cell growth inside microfluidic devices under flow and static conditions. Panel of phase contrast images of caco-2 cells at day 2, day 4 and day 7 of growth inside microfluidic channels under constant flow that exerts a shear stress of 0.02 dynes/cm^2^ to the cells and static conditions with media change every day. The arrowheads point out the villi-like structures formed by the cells inside the microfluidic device (scale bars = 200 µm).

Following the 7-day culture period under flow conditions, the microfluidic device was carefully divided into two halves to prepare them for different imaging modalities. One half was processed for scanning electron microscopy (SEM), where images revealed the presence of villi-like structures, consistent with in-vivo architecture (Figure 5A-B). The second half of the device was fixed and subsequently immunostained for analysis by confocal microscopy. Obtained Z-stack (Figure 5C) showed the presence of the three-dimensional structures of the Caco-2 cell layer subjected to shear stress across the microchannel. Orthogonal views of the z-stack (Figure 5D-E) further revealed the spatial arrangement and vertical profile of the cell monolayer. The height of the highest structure was measured to be 330 µm, it is important to note that this measurement may underestimate the true height, as the upper limit of the microscope’s Z-axis was reached. Confocal imaging also showed an F-actin distribution along the border of the structures (Figure 5B), indicative of polarized epithelial cells, and the presence of mucin production in the intercellular space, characteristic of differentiated mucus-producing cells (Figure 5F).

**Figure 5.**
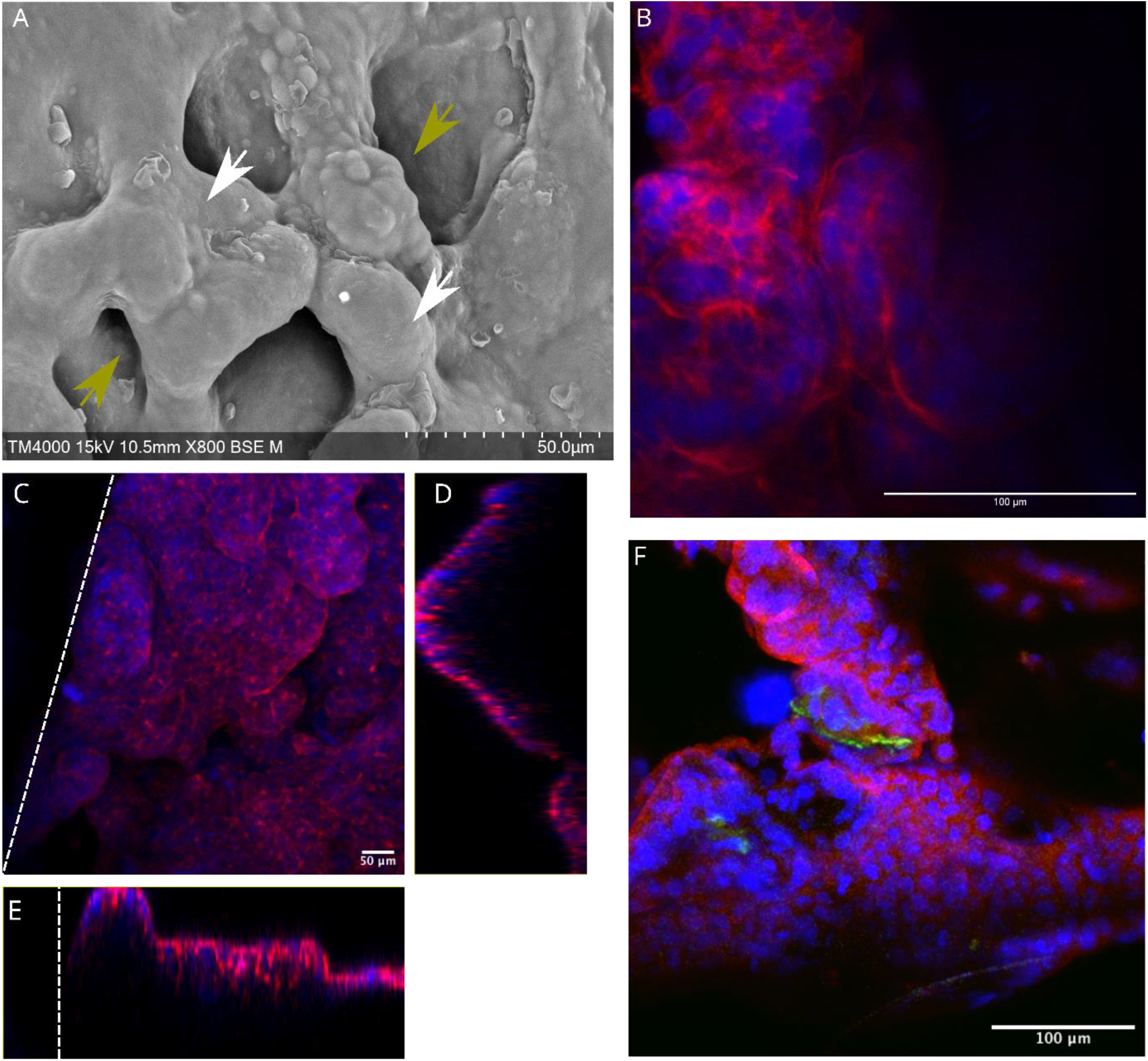
Morphological characterization of differentiated caco-2 cell monolayer after 7 days of cell culture under flow. A. Scanning electron microscope (SEM) imaging showing villi-like structures (white arrowhead) and crypt-like invaginations (green arrowhead). B and C. Z stack of villi-like structures immunostained to show villin (red) and nuclei (blue). D and E. Orthogonal view at highest point of the area (dotted line represents the microchannel wall). F. Z stack of differentiated epithelium showing the production of mucin by the cells (MUC-2 in green), F-actin in red and nuclei (DAPI in blue).

Once the presence of villi-like structures inside the microfluidic device was confirmed, the cellular layer was infected with *Candida albicans* yeast in the exponential growth phase. The infection was allowed to progress for 24 hours, with observations made at different stages. Hyphal development was first observed 3 hours after the yeast adhered to the intestinal epithelium. At the 24-hour mark, the cells and yeast were fixed and prepared for SEM and confocal microscopy. *C. albicans* employs two well-documented mechanisms for epithelial invasion: epithelial-driven endocytosis (also known as induced endocytosis) occurring in the initial hours of infection, and active penetration, which becomes prominent during later stages of interaction but there has been observation of other routes that could be taken by the yeast as the paracellular route that alters the adherent junctions and tight junctions of cells (40). SEM observation of the infected samples showed filaments of *Candida albican*s piercing the cellular membrane of the Caco-2 cells (Figure 6A) as well as the disruption of the tight junctions in order to cross the cellular layer (Figure 6B). Confocal observation showed the yeast filaments extending into the interior of the Caco-2 cells and the accumulation of actin in the place of piercing, an event to be known to happen during induced endocytosis (41) (Figure 6C).

**Figure 6.**
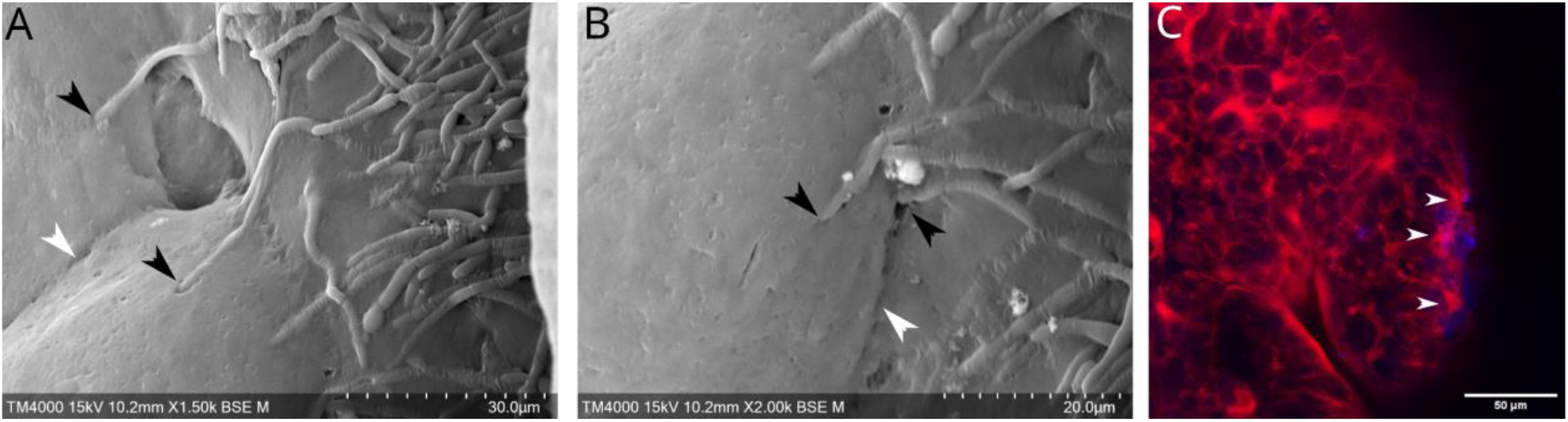
Morphological characterization of differentiated intestinal epithelium after *Candida albicans* infection. A and B. Scanning electron microscope (SEM) imaging of intestinal epithelium after 24 hours of *Candida albicans* infection: white arrowhead shows tight junctions between cells, black arrowhead shows hyphae perforating cellular membrane C. Z stack image of differentiated intestinal epithelium (Caco-2 F-actin in red) infected by *Candida albicans* (yeast stained with Calcofluor White, blue).

To determine whether *C. albicans* infection affected the barrier function of the epithelial cell layer, we conducted a permeability assay by perfusing cRPMI-PR medium with a high concentration of fluorescein (332 kDa) in the upper channel and measuring the fluorescent intensity of the diffused molecules in the lower channel to obtain the apparent permeability (Figure 7A). The naked membrane exhibited an apparent permeability of 2.5 × 10^−5^ cm/s, consistent with literature values for other collagen membranes tested with fluorescein as the transported molecule (43). Upon formation of the epithelial cell layer, as expected, the permeability decreased by one order of magnitude compared to the naked membrane, this value of 1.5 × 10^−6^ cm/s is in good agreement with the results reported in a gut-on-chip system after 5 days of cell growth (37), validating the establishment of a functional intestinal epithelial barrier within our system. In contrast, the infected layer showed a higher apparent permeability among the three conditions, with a value of 1.5 × 10^−4^ cm/s. This increase can likely be attributed to the fact that *Candida* hyphae not only damaged the epithelium but also compromised the collagen membrane, facilitating significant diffusion into the lower channel. Finally, the comparison of thigh junctions between a healthy epithelial layer and an infected epithelial layer was done as they have a major role on the barrier function of the intestinal epithelium. Figure 7B shows the continuous network of tight junctions present between caco-2 cells while figure 7C shows the lack of tight junctions in a great part of the layer, showing the impact of the infection over the cells.

**Figure 7.**
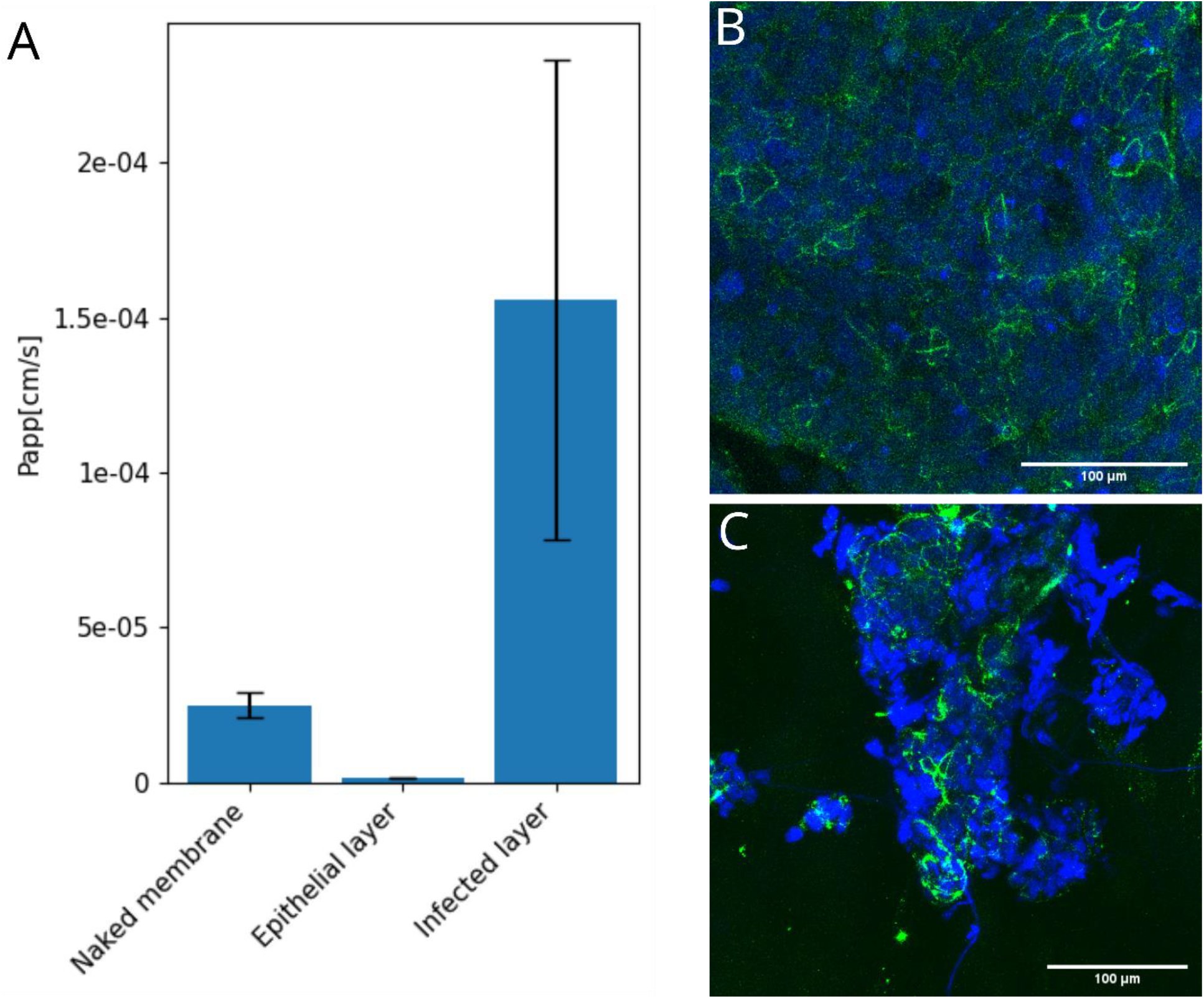
Permeability assays. **A**. Apparent permeability comparison of collagen type I membrane without cells, with differentiated epithelial layer and epithelial layer infected or not by *Candida albicans* yeast. B. Healthy epithelial layer showing tight junctions (ZO-1 in green) between caco-2 cells, and nuclei (DAPI in blue) C. Infected epithelial layer showing damaged tight junctions by *C. albicans* hyphae (Calcofluor White and DAPI in blue).

## Conclusions

In this study, a two-chamber gut-on-chip was fabricated incorporating a biocompatible collagen-type I membrane, to mimic the natural ECM and get closer to the environment intestinal cells face *in vivo*. Caco-2 cells adhered to the membrane, formed a confluent monolayer and flow boosted their differentiation into villi-like structures. The presence of tight junctions, mucin production, and barrier function indicated successful differentiation of the epithelium.

We also investigated the device’s capacity to model host-pathogen interactions by infecting the cell layer with *Candida albicans*. We observed hyphal invasion of the epithelial cells, demonstrating the device’s capacity to model infection processes and resulting changes in barrier function, specifically increased layer permeability. These results have significant implications for the development of more physiologically relevant collagen-based *in vitro* models for drug testing, disease modeling, and studying host-pathogen interactions at the intestinal barrier.

## Acknowledgments

We thank Armelle Ménard (Bordeaux Institute of Oncology, Bordeaux University) for kindly providing the Caco-2 cell line, Lamia Azzi-Martin for her advice for immunostaining protocols and Melanie Bonhivers (Fundamental Microbiology and Pathogenicity Lab, Bordeaux University) for generously proving the secondary antibodies for confocal observations. This study received financial support from the French government in the framework of the University of Bordeaux’s France 2030 program/GPR LIGHT and the New Models in Oncology Impulsion Research Network (RRI NewMoon).

## Notes

### Competing Interest Statement

The authors have declared no competing interest.

## References

1. Jing B, Wang ZA, Zhang C, Deng Q, Wei J, Luo Y, et al. Establishment and Application of Peristaltic Human Gut-Vessel Microsystem for Studying Host–Microbial Interaction. Front Bioeng Biotechnol. 2020;8:272. 10.3389/fbioe.2020.00272

2. Xiang Y, Wen H, Yu Y, Li M, Fu X, Huang S. Gut-on-chip: Recreating human intestine in vitro. J Tissue Eng. 2020;11:204173142096531. 10.1177/2041731420965318

3. Wang L, Murthy SK, Fowle WH, Barabino GA, Carrier RL. Influence of micro-well biomimetic topography on intestinal epithelial Caco-2 cell phenotype. Biomaterials. 2009;30(36):6825– 34. 10.1016/j.biomaterials.2009.08.046

4. Wang L, Murthy SK, Barabino GA, Carrier RL. Synergic effects of crypt-like topography and ECM proteins on intestinal cell behavior in collagen based membranes. Biomaterials. 2010;31(29):7586–98. 10.1016/j.biomaterials.2010.06.036

5. Grassart A, Malardé V, Gobaa S, Sartori-Rupp A, Kerns J, Karalis K, et al. Bioengineered Human Organ-on-Chip Reveals Intestinal Microenvironment and Mechanical Forces Impacting Shigella Infection. Cell Host & Microbe. 2019;26(3):435-444.e4. 10.1016/j.chom.2019.08.007

6. Gérémie L, Ilker E, Bernheim-Dennery M, Cavaniol C, Viovy J-L, Vignjevic DM, et al. Evolution of a confluent gut epithelium under on-chip cyclic stretching. Phys Rev Research. 2022;4(2):023032. 10.1103/PhysRevResearch.4.023032

7. Thomas DP, Zhang J, Nguyen N-T, Ta HT. Microfluidic Gut-on-a-Chip: Fundamentals and Challenges. Biosensors. 2023;13(1):136. 10.3390/bios13010136

8. Bhatia SN, Ingber DE. Microfluidic organs-on-chips. Nat Biotechnol. 2014;32(8):760–72. 10.1038/nbt.2989

9. Fedi A, Vitale C, Ponschin G, Ayehunie S, Fato M, Scaglione S. In vitro models replicating the human intestinal epithelium for absorption and metabolism studies: A systematic review. Journal of Controlled Release. 2021;335:247–68. 10.1016/j.jconrel.2021.05.028

10. Pimenta J, Ribeiro R, Almeida R, Costa PF, Da Silva MA, Pereira B. Organ-on-Chip Approaches for Intestinal 3D In Vitro Modeling. Cellular and Molecular Gastroenterology and Hepatology. 2022;13(2):351–67. 10.1016/j.jcmgh.2021.08.015

11. Nikolaev M, Mitrofanova O, Broguiere N, Geraldo S, Dutta D, Tabata Y, et al. Homeostatic mini-intestines through scaffold-guided organoid morphogenesis. Nature. 2020;585(7826):574–8. 10.1038/s41586-020-2724-8

12. Gjorevski N, Nikolaev M, Brown TE, Mitrofanova O, Brandenberg N, DelRio FW, et al. Tissue geometry drives deterministic organoid patterning. Science. 2022;375(6576):eaaw9021. 10.1126/science.aaw9021

13. Verhulsel M, Simon A, Bernheim-Dennery M, Gannavarapu VR, Gérémie L, Ferraro D, et al. Developing an advanced gut on chip model enabling the study of epithelial cell/fibroblast interactions. Lab Chip. 2021;21(2):365–77. 10.1039/D0LC00672F

14. Mitrofanova O, Nikolaev M, Xu Q, Broguiere N, Cubela I, Camp JG, et al. Bioengineered human colon organoids with in vivo-like cellular complexity and function. Cell Stem Cell. 2024;31(8):1175-1186.e7. 10.1016/j.stem.2024.05.007

15. Jalili-Firoozinezhad S, Prantil-Baun R, Jiang A, Potla R, Mammoto T, Weaver JC, et al. Modeling radiation injury-induced cell death and countermeasure drug responses in a human Gut-on-a-Chip. Cell Death Dis. 2018;9(2):223. 10.1038/s41419-018-0304-8

16. Jalili-Firoozinezhad S, Gazzaniga FS, Calamari EL, Camacho DM, Fadel CW, Bein A, et al. A complex human gut microbiome cultured in an anaerobic intestine-on-a-chip. Nat Biomed Eng. 2019;3(7):520–31. 10.1038/s41551-019-0397-0

17. Pasman, Thijs, et al. Flat and Microstructured Polymeric Membranes in Organs-on-Chips. Journal of The Royal Society Interface. 2018;15(144):20180351. 10.1098/rsif.2018.0351

18. Shin W, Kim HJ. 3D in vitro morphogenesis of human intestinal epithelium in a gut-on-a-chip or a hybrid chip with a cell culture insert. Nat Protoc. 2022;17(3):910–39. 10.1038/s41596-021-00674-3

19. Rahimnejad, Maedeh, et al. Engineered Biomimetic Membranes for Organ-on-a-Chip. ACS Biomaterials Science & Engineering. 2022. 10.1021/acsbiomaterials.2c00531

20. Pérez-González C, Ceada G, Greco F, Matejčic M, Gómez-González M, Castro N, et al. Mechanical compartmentalization of the intestinal organoid enables crypt folding and collective cell migration. Nat Cell Biol. 2021;23(7):745–57. 10.1038/s41556-021-00699-6

21. Mondrinos MJ, Yi Y-S, Wu N-K, Ding X, Huh D. Native extracellular matrix-derived semipermeable, optically transparent, and inexpensive membrane inserts for microfluidic cell culture. Lab Chip. 2017;17(18):3146–58. 10.1039/C7LC00317J

22. Pompili S, Latella G, Gaudio E, Sferra R, Vetuschi A. The Charming World of the Extracellular Matrix: A Dynamic and Protective Network of the Intestinal Wall. Front Med. 2021;8:610189. 10.3389/fmed.2021.610189

23. Meran, Laween, et al. Intestinal Stem Cell Niche: The Extracellular Matrix and Cellular Components. Stem Cells International. 2017;2017:7970385. 10.1155/2017/7970385

24. Graham, Martin F., et al. Collagen Content and Types in the Intestinal Strictures of Crohn’s Disease. Gastroenterology. 1988;94(2):257–65. 10.1016/0016-5085(88)90411-8

25. Lee JS, Romero R, Han YM, Kim HC, Kim CJ, Hong J-S, et al. Placenta-on-a-chip: a novel platform to study the biology of the human placenta. The Journal of Maternal-Fetal & Neonatal Medicine. 2016;29(7):1046–54. 10.3109/14767058.2015.1038518

26. Takezawa T, Ozaki K, Nitani A, Takabayashi C, Shimo-Oka T. Collagen Vitrigel: A Novel Scaffold that can Facilitate a Three-Dimensional Culture for Reconstructing Organoids. Cell Transplant. 2004;13(4):463–74. 10.3727/000000004783983882

27. Corral-Nájera K, Chauhan G, Serna-Saldívar SO, Martínez-Chapa SO, Aeinehvand MM. Polymeric and biological membranes for organ-on-a-chip devices. Microsyst Nanoeng. 2023;9(1):107. 10.1038/s41378-023-00579-z

28. Salimbeigi G, Vrana NE, Ghaemmaghami AM, Huri PY, McGuinness GB. Basement membrane properties and their recapitulation in organ-on-chip applications. Materials Today Bio. 2022;15:100301. 10.1016/j.mtbio.2022.100301

29. Tasiopoulos CP, Gustafsson L, Van Der Wijngaart W, Hedhammar M. Fibrillar Nanomembranes of Recombinant Spider Silk Protein Support Cell Co-culture in an In Vitro Blood Vessel Wall Model. ACS Biomater Sci Eng. 2021;7(7):3332–9. 10.1021/acsbiomaterials.1c00612

30. Youn J, Hong H, Shin W, Kim D, Kim HJ, Kim DS. Thin and stretchable extracellular matrix (ECM) membrane reinforced by nanofiber scaffolds for developing in vitro barrier models. Biofabrication. 2022;14(2):025010. 10.1088/1758-5090/ac4dd7

31. Zamprogno P, Wüthrich S, Achenbach S, Thoma G, Stucki JD, Hobi N, et al. Second-generation lung-on-a-chip with an array of stretchable alveoli made with a biological membrane. Commun Biol. 2021;4(1):168. 10.1038/s42003-021-01695-0

32. Wang C, Tanataweethum N, Karnik S, Bhushan A. Novel Microfluidic Colon with an Extracellular Matrix Membrane. ACS Biomater Sci Eng. 2018;4(4):1377–85. 10.1021/acsbiomaterials.7b00883

33. Vilardi A, Przyborski S, Mobbs C, Rufini A, Tufarelli C. Current understanding of the interplay between extracellular matrix remodelling and gut permeability in health and disease. Cell Death Discov. 2024;10(1):258. 10.1038/s41420-024-02015-1

34. Asal M, Rep M, Bontkes HJ, Van Vliet SJ, Mebius RE, Gibbs S. Towards Full Thickness Small Intestinal Models: Incorporation of Stromal Cells. Tissue Eng Regen Med. 2024;21(3):369–77. 10.1007/s13770-023-00600-6

35. Hojat, N., Gentile, P., Ferreira, A.M. et al. Automatic pore size measurements from scanning electron microscopy images of porous scaffolds. J Porous Mater 30, 93–101 (2023). 10.1007/s10934-022-01309-y

36. Kim HJ, Huh D, Hamilton G, Ingber DE. Human gut-on-a-chip inhabited by microbial flora that experiences intestinal peristalsis-like motions and flow. Lab Chip. 2012;12(12):2165. 10.1039/c2lc40074j

37. Huh D, Kim HJ, Fraser JP, Shea DE, Khan M, Bahinski A, et al. Microfabrication of human organs-on-chips. Nat Protoc. 2013;8(11):2135–57. 10.1038/nprot.2013.133

38. Kim HJ, Ingber DE. Gut-on-a-Chip microenvironment induces human intestinal cells to undergo villus differentiation. Integr Biol. 2013;5(9):1130. 10.1039/c3ib40126j

39. Basmaciyan L, Bon F, Paradis T, Lapaquette P, Dalle F. Candida Albicans Interactions With The Host: Crossing The Intestinal Epithelial Barrier. Tissue Barriers. 2019;7(2):e1612661. 10.1080/21688370.2019.1612661

40. Wells JM, Rossi O, Meijerink M, Fransen F. Epithelial crosstalk at the microbiota–gut interface. Gut Microbes. 2010;1(3):123–30. 10.1111/j.1462-5822.2009.01394.x

41. Zhang Y, Zhou Y, Chen X, Zhang Y, Yuan Y, Zhao Y, et al. Bioinspired Hydrogel Membranes with Tunable Permeability for Gut-on-a-Chip Applications. ACS Biomater Sci Eng. 2020;6(5):2918–29. 10.1021/acsbiomaterials.0c00297

